# An Optimally Weighted Combination Method to Detect Novel Disease Associated Genes Using Publicly Available GWAS Summary Data

**DOI:** 10.1101/709808

**Authors:** Jianjun Zhang, Samantha Gonzales, Jianguo Liu, Xiaoyi Raymond Gao, Xuexia Wang

**Affiliations:** Department of Mathematics, University of North Texas,1155 Union Circle #311430 Denton, TX 76203-5017; Department of Computer Science and Engineering, University of North Texas, Discovery Park 3940 N. Elm, Denton, TX 76203-5017; Department of Ophthalmology and Visual Science, The Ohio State University, Columbus, OH 43212, USA; Department of Biomedical Informatics, The Ohio State University, Columbus, OH 43212, USA; Division of Human Genetics, The Ohio State University, Columbus, OH 43212, USA

## Abstract

Gene-based analyses offer a useful alternative and complement to the usual single nucleotide polymorphism (SNP) based analysis for genome-wide association studies (GWASs). Using appropriate weights (pre-specified or eQTL-derived) can boost statistical power, especially for detecting weak associations between a gene and a trait. Because the sparsity level or association directions of the underlying association patterns in real data are often unknown and access to individual-level data is limited, we propose an optimal weighted combination (OWC) test applicable to summary statistics from GWAS. This method includes burden tests, weighted sum of squared score (SSU), weighted sum statistic (WSS), and the score test as its special cases. We analytically prove that aggregating the variants in one gene is the same as using the weighted combination of Z-scores for each variant based on the score test method. We also numerically illustrate that our proposed test outperforms several existing comparable methods via simulation studies. Lastly, we utilize schizophrenia GWAS data and a fasting glucose GWAS meta-analysis data to demonstrate that our method outperforms the existing methods in real data analyses. Our proposed test is implemented in the R program OWC, which is freely and publicly available.

## Introduction

Even though genome-wide association studies (GWASs) have been remarkably successful in identifying a large number of genetic variants associated with complex traits and diseases, these identified variants can only explain a small to modest fraction of the heritability (Manolio et al., 2009). Larger sample sizes and more powerful statistical tests are needed to boost power, especially for weakly associated variants with small effect sizes or low frequency variants. Due to various reasons, it is often difficult for researchers to obtain access to individual level data, and thus difficult to obtain a sufficient sample size for reliable analysis. The increasingly public availability of genome-wide association study (GWAS) summary statistics, e.g. p-values, effect sizes, directions of effects, or estimated statistics for individual single nucleotide polymorphisms (SNPs) motivates us to develop methods for further analyzing GWAS by integrating the results from standard single variant analysis using GWAS summary data. Methods based on summary statistics can also be viewed as a complementary approach to the traditional single variant single trait association test.

The power of different gene-based tests depends on the underlying genetic architecture, which can differ in number, effect size, and effect direction of the causal variants in different genes. The proposed weighted combination method is a general and flexible method that performs significantly better when testing for weak association due to small effect sizes or low frequency variants. When testing for genetic associations, a proper choice of weights can boost power substantively in gene-based analysis. For instance, the burden test (Li and Leal, 2008; Morgenthaler and Thilly, 2007) and weighted sum of squared score (SSU) test (Pan 2009) can be viewed as weighted combination methods, where the burden test and SSU test set the same weight and use the Z-score as a weight for each variant, respectively. A statistical challenge is that, due to unknown true association patterns, there is no uniformly most powerful test to detect single trait associated genes; an association test may perform well for one dataset, but not necessarily for another (Kwak and Pan 2015). For example, the presence of non-associated SNPs will largely diminish the power of a standard test if no effective SNP selection or weighting is adopted (Petersen et al., 2013). Additionally, SSU provides a robust test that is particularly powerful in the presence of protective, deleterious, and null variants, but is less powerful than the burden test when a large number of variants in a region are causal and in the same direction. In the Method section of this paper, we show that using a score test to test the aggregated variants in a gene is the same as using the weighted combination of Z-scores for the variants in the considered region. Thus, existing popular weighted combination methods based on individual genotype and phenotype data can be modified and implemented based on summary data. In this paper, we propose a new gene based genetic association test method, the optimal weighted combination (OWC) test, which can use GWAS summary statistics as input. More importantly, OWC includes the burden test, SSU, WSS, and score test as its special cases, and OWC is an optimally weighted combination test that can reach maximized power.

To evaluate the performance of the proposed method, we conducted extensive simulation studies and real data analysis. We compared our method OWC with five existing gene-based methods: gene-based association test that uses extended Simes procedure (GATES) (Li et al., 2011), adaptive sum of powered score tests (aSPU) (Kwak and Pan 2015), and three methods proposed by Guo and Wu (2018a): sum test (ST), squared sum test (S2T), adaptive test (AT). All methods are designed for single trait association study. GATES adopts an extended Simes procedure to correct multiple testing issues while calculating the p-value quickly based on SNP summary statistics. The aSPU method estimates and selects the most powerful test among a class of so-called sum of powered score (SPU) tests. ST is a type of burden test statistic (Madsen and Browning, 2009), S2T is a type of SKAT statistic (Wu et al., 2011) and AT is equivalent to the SKAT-O statistic (Lee, Wu and Lin 2012). Simulation results demonstrate that our proposed method OWC outperforms the five comparable methods. Application to the schizophrenia GWAS summary data obtained from the Psychiatric Genomics Consortium (PGC) and fasting glucose GWAS meta-analysis summary data obtained from the UK Biobank component of the European DIAMANTE study indicate that our method performs better than existing methods.

## Material and Methods

### Inference

Consider a raw data set of a sample including *n* individuals, where each individual has been genotyped at M variants in a genomic region (gene or pathway). Denote *y*_*i*_ as the trait value of the *i^th^* individual for either a quantitative or qualitative trait (1 for cases and 0 for controls for a qualitative trait) and denote *X*_*i*_ = (*x_i1_*, …, *x*_*iM*_)′ as the genotypic score of the *i^th^* individual, where *x*_*im*_ ∈ {0, 1, 2} is the number of minor alleles that the *i^th^* individual has at the *m^th^* variant. *x*_*im*_ can also be imputated doseage or the number of minor alleles in dominant or recessive coding of genotypes that the *i^th^* individual has at the *m^th^* variant.

We use the generalized linear model to model the relationship between the trait (quantitative or qualitative trait) and the genetic variants in the considered region:

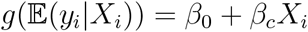

where *g*(*⋅*) is a monotone “link” function and *β_c_* is the parameter of interest. Testing the association of the genetic variants in the considered region is equivalent testing the effect of the weighted combination of variants 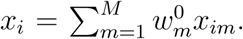. Under the generalized linear model, we can use the score test statsitic to test the null hypothesis *H*_0_: *β_c_* = 0, which is given by

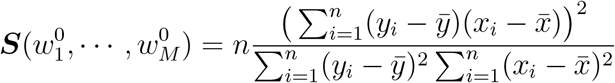

where the score test statistic ***S*** can be viewed as a function of weight 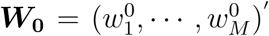. Let ***X*** = (*X*_1_, …, *X*_*n*_)′, *Y* = (*y*_1_, …, *y*_*n*_)′ and 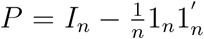 where 1_*n*_ represents a column vector containing all ones. Then, we have 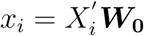. We can rewrite the score test as:

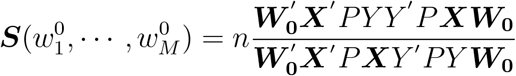

Derivation of the aforementioned score test can be found in the Appendix section. To test the association between a single trait and a single variant, a Z test is generally employed. To test the main effect of the *m^th^* variant in the considered region, we use the Z test: 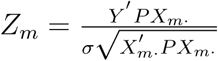 where 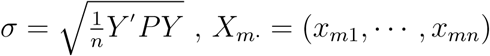. Let LD matrix ***R*** = *diag*(***D***)^*−*1*/*2^***D**diag*(***D***)^*−*1*/*2^ where ***D*** = ***X**′ P **X*** and *diag*(***D***) denote the diagonal matrix of ***D***. When GWAS summary statistics such as the Z-statistics and the LD matrix for SNP-SNP correlations are available, the score test can be written as:

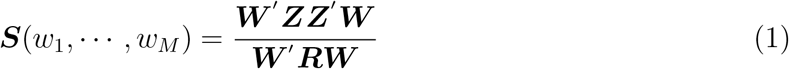

where ***Z*** = (*Z*_1_, …, *Z*_*M*_)′ and ***W*** = (*w*_1_, …, *w*_*M*_)′ = *diag*(***D***)^1*/*2^***W***_0_ (see Appendix). From Equation (1), the score test statistic ***S*** is equivalent to a linear weighted test statistic based on Z-scores:

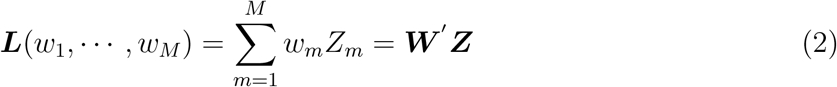

where ***Z*** follows multivariate normal distribution with mean **0** and covariance matrix ***R*** under null hypothesis (Zhang et al. 2018). This conclusion clearly demonstrates that testing the weighted combination of variants in a considered region using the score test is the same as using the weighted combination of Z-scores for those variants.

### Gene-based tests

The true value of the aforementioned weight function ***W*** = (*w*_1_, …, *w*_*M*_)′ is unknown and must be determined biologically or empirically. Therefore, in real data analysis, we should give reasonable values of weights in advance for a gene-based test. If all or most of the variants have nearly equal effect size in the same direction of association, we set *w*_*m*_ = 1 for *m* = 1, …, *M*, and the test becomes the burden test ***L***_*B*_ = ***L***(1, …, 1), which sums up the association signals across all the variants and obtains high power. If we believe that the causal SNPs would be subject to “purifying selection” and thus appear less frequently in the population than neutral SNPs, we set 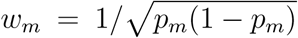, where *p*_*m*_ denotes the minor allele frequency (MAF) of the *m^th^* variant, and obtain 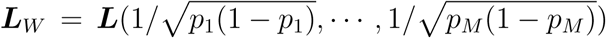, which is the weighted sum statistic (WSS) (Madsen and Browning, 2009). If we assume that the values of the weights ***W*** come from gene expression or functional annotation data, the test degenerates into the PathSPU(1) test (Wu and Pan 2018). We know that ***S***(*w*_1_, …, *w*_*M*_) follows central chi-square distribution with 1 degree of freedom 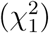 and ***L***(*w*_1_, …, *w*_*M*_) follows multivariate normal distribution with mean **0** and covariance matrix ***W***′***RW*** under the null hypothesis when the choice of the weight function ***W*** is not proportional to ***Z***.

As a function of ***W*** = (*w*_1_, …, *w*_*M*_)′, we can also choose an optimal ***W*** that maximizes either the score test ***S***(*w*_1_, …, *w*_*M*_) or the linear weighted test statistic ***L***(*w*_1_, …, *w*_*M*_). In particular (Li and Lagakos 2006), we have

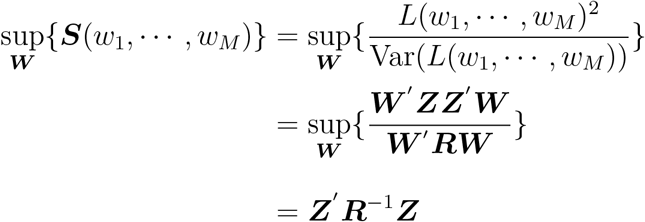

When 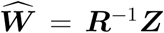, the score test statistic ***S***(*w*_1_, …, *w*_*M*_) reaches its maximum. Based on the asymptotic null distribution of ***Z*** in Equation (2), we define the score test 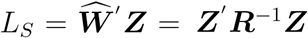 which follows central chi-square distribution with *M* degrees of freedom 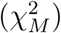. The optimal weights can be obtained when the linear weighted test statistic reaches its maximum (Basu and Pan 2011). If we consider the correlation matrix ***R*** as a diagonal matrix ***A*** = *diag*(*a*_1_, …, *a*_*M*_), ***R*** = ***A***, where 0 *< a_i_ ≤* 1, then ***W*** = ***A**^−^*^1^***Z***. The score test in Equation (1) or the linear weighted test in Equation (2) will reach its maximum when ***W*** = ***A**^−^*^1^***Z***. To test the association between SNPs in a considered region and a trait, Kwak and Pan (2015) proposed a class of sum of powered score (SPU) tests, along with its data-adaptive version (aSPU), 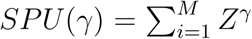, and *γ* = 1, 2, …, 8*, ∞*. The SPU method can also be viewed as a special weighted combination test method with weight ***W*** = ***Z**^γ−^*^1^ and aSPU can be viewed as an data-adaptive weighted combination test method.

When the diagonal matrix *A* is the identity matrix: ***A*** = ***I***, we denote the test in Equation (2) as *L*_*Q*_ = ***Z**′**Z***, which is the same as the sum of squared score test (SSU) (Pan 2011) or the variance component test (Tzeng et al. 2011). Based on the asymptotic null distribution of ***Z*** in Equation (2), the test *L*_*Q*_ = ***Z**′**Z*** follows a mixture of chi square distribution under the null hypothesis: 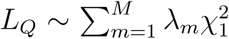, where *λ*_1_, …, *λ_M_* are the eigenvalues of ***R***. Particularly, if we set the diagonal element of ***A*** as the beta distribution density function with pre-specified shape parameters as 1 and 25 evaluated at the corresponding sample MAF in the data, it degenerates into the sequence kernel association test (SKAT) for rare variants (Wu et al., 2011). If the value of the diagonal elements of ***A*** comes from a set of gene expression derived weights, it degenerates into PathSPU(2) test method (Wu and Pan, 2018). Naturally, these two methods (SKAT and PathSPU(2)) all follow a mixture of chi square distribution under the null hypothesis. In our paper, we only consider GWAS summary data for common variants, thus we set ***A*** as the identity matrix for this case.

To combine the strength of *L_B_, L_W_, L_S_*, and *L*_*Q*_, we consider their weighted average:

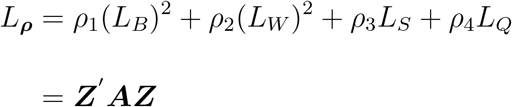

where ***A*** = *ρ*_1_**11**′+ *ρ*_2_***W W***′+ *ρ*_3_***R**^−^*^1^ + *ρ*_4_***I***, **1** denotes a column vector containing all 1s, *ρ*_1_ + *ρ*_2_ + *ρ*_3_ + *ρ*_4_ = 1, and 0 *≤ ρ_i_ ≤* 1 for *i* = 1, 2, 3, 4. For a fixed ***ρ***, *L*_**ρ**_ is distributed as a linear combination of independent central 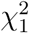 random variables under the null hypothesis:

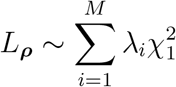

where 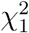 denotes a central *χ*^2^ random variable with 1 degrees of freedom and *λ_i_* for *i* = 1, …, *M* are the eigenvalues of ***RA*** (Pan 2009). We propose a novel method which reaches the optimal weighted combination (OWC) for a set of values of ***ρ*** using the minimum p-value across the values of ***ρ***

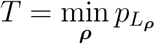

where 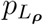 is the p-value computed based on *L_**ρ**_*. Naturally, *T* can be obtained by a simple grid search across a range of ***ρ***: set a grid {*ρ*_1_, *ρ*_2_, *ρ*_3_, *ρ*_4_*}*, then the test statistic 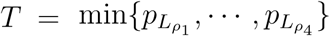. In practice, we search over *ρ_i_ ∈ (*0, 0.1, 0.2, 0.3, 0.4, 0.5, 0.6, 0.7, 0.8, 0.9, 1) for *i* = 1, 2, 3, 4. Specifically, if *ρ*_2_ = *ρ*_3_ = 0 in ***ρ***, *L*_**ρ**_ can be rewritten as *ρ*_1_(*L*_*B*_)^2^ + (1 *− ρ*_1_)*L*_*Q*_, which is equivalent to SKAT-O test method (Lee, Wu and Lin 2012).

### P-value estimation

Monte Carlo simulations are used to obtain the p-values for *T* in a single layer of simulations. Briefly, after obtaining ***R***, we first simulate null scores of ***Z***^(*b*)^ ∼ *N* (0*, **R***) for *b* = 1, …, *B*.

Then, we use the null scores to calculate the null test statistic *T ^b^* following the aforementioned procedure for each b, and then the p-value of the test is the proportion of the number of the null test statistic *T ^b^* with *T ^b^ ≤ T* (Kwak and Pan, 2016). A larger B is needed to estimate a smaller p-value.

The aforementioned vector ***Z***^(*b*)^ can be generated in the following way (Zhou, 2014): we first generate a vector ***L*** with *M* elements where each element is independently generated from a standard univariate normal distribution with mean 0 and variance 1; that is, ***L** ∼ N* (**0***, **I***). We then have ***Z***^(*b*)^ = ***DL***, where ***D*** is obtained from Cholesky decompositions of ***R*** with ***R*** = ***DD***′. Specifically, for the test statistic *T* (***Z**, **R***) as a function of ***Z*** and ***R***, we can estimate its p-value in detail as follows:

1. Generate independent ***Z***^(*b*)^ *∼ N* (0*, **R***) for *b* = 1, …, *B*.
2. Using asymptotic distribution of *L_**ρ**_* under null hypothesis, calculate the null test statistic *T* by searching across a range of ***ρ*** for ***Z*** and ***Z***^(*b*)^, respectively.
3. Finally, the p-value for the *T* test, *p*_*T*_, the p-value of *T* (***Z**, **R***) is

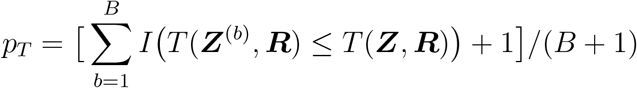

If the Z statistic in the summary data is not provided, we need to first transform the p-value in the summary data into a Z statistic using *Z* = sign(*β*)Φ^*−*1^(1 *− p/*2), where Φ is the cumulative distribution function of the standard univariate normal distribution. Then, a similar procedure can be used to obtain the p-value of the test *T*.

### Comparison of methods

We compared the performance of our proposed method (OWC) with five aforementioned methods: 1) three methods proposed by Guo and Wu (2018a): sum test (ST), squared sum test (S2T), adaptive test (AT), 2) a method proposed by Kwak and Pan (2015): adaptive sum of powered score tests (aSPU), and 3) a method proposed by Li et al. (2011): gene-based association test that uses extended Simes procedure (GATES). We briefly introduce each of the methods as follows:

1. Sum test (ST), 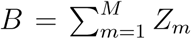, a type of burden test statistic (Madsen and Browning, 2009).
2. Squared sum test (S2T), 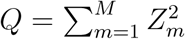, a type of SKAT statistic (Wu et al., 2010) and equivalent to the weighted sum of squared score (SSU) test (Pan 2009).
3. Adaptive test (AT), *T* = min_*ρ∈*[0,1]_ *P* (*Q_ρ_*), where *Q_ρ_* = (1 *− ρ*)*Q* + *ρB*^2^ and *P* (*Q_ρ_*) denotes the corresponding p-value.
4. Adaptive sum of powered score tests (aSPU), aSPUs=min_*γ∈*Γ_ *P*_*SPUs*(*γ*)_ where 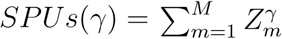.
5. Gene-based association test that uses extended Simes procedure (GATES), 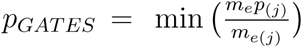 where *m*_e_ is the effective number of independent p-values among the M SNPs, *p*_(*j*)_ is the *j^th^* smallest p-value and *m*_*e*(*j*)_ is the effective number of independent p-values among the top *j* SNPs.

Because ***Z*** ∼ MVN(**0**, ***R***), 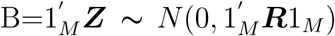 where 1_*M*_ denotes a column vector of length *M* in which elements are all 1’s, 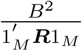 follows *χ*1 distribution. Q=***Z***73×2032;***Z*** is asymptotically distributed as the weighted sum of independent *χ*_1_ random variables with the weights being the eigenvalues of ***R***. The p-value of *T* can be efficiently and accurately computed by a one-dimensional numerical integration where we will search over *ρ ∈* (0, 0.01, 0.04, 0.09, 0.16, 0.25, 0.5, 1) following Wu et al. (2016). The p-value of the three statistics can be obtained using the “sats” function in the “mkatr” package in R. For adaptive sum of powered score tests (aSPU), Monte Carlo simulations are used to obtain the p-values, and this test can be obtained using the “aSPUs” function in the “aSPU” package in R. To estimate the p-value of the gene-based association test that uses extended Simes procedure (GATES), we use the “gates” function in the “COMBAT” package in R.

## Results

### Simulations Studies

To evaluate the performance of the proposed method OWC, we conducted extensive simulation studies to compare the type I error and power of OWC with the five comparable methods following the simulation setting in Guo and Wu (2018b). The LD between SNPs are estimated using the 1000 Genomes project (1000 Genomes Project Consortium, 2012) because Kwak and Pan (2015) have shown that the performance using any reference data from the same ancestry in estimating LD among SNPs is mostly satisfactory with an estimated inflation factor close to 1.

### Simulation of the Type I Error

For the type I error evaluation, instead of generating genotype and phenotype data, we simulate the test statistic ***Z*** from a multivariate normal distribution *N* (**0***, **R***) where ***R*** denotes the corresponding LD matrix of the gene *EP B*41 (erythrocyte membrane protein band 4.1) which is used in our simulation studies. Gene *EP B*41 colocalizes with *AMP A* receptors at excitatory synapses and is thought to mediate the interaction of the *AMP A* receptors with the cytoskeleton (Shen et al., 2000). Numerous studies have demonstrated brain region- and subunit-specific abnormalities in the expression of subunits of the AMPA subtype of glutamate receptors in schizophrenia (Tucholski et al., 2013). We consider four different significance levels: *α* = 10^*−*3^, 10^*−*4^, 10^*−*5^, and 2.80 *×* 10^*−*6^. The estimated type I error rates are summarized in Table 1. In the simulation, p-values of our method and aSPU are estimated by performing 10^6^ times permutations and type I error rates are calculated based on 10^6^ replicates. From this table, we can see that the type I error rates of all of the methods are well controlled. Thus, all the tests are valid.

**Table 1:** Estimated type I error rates for different test methods.

| α-level | ST | S2T | AT | GATES | aSPU | OWC |
| --- | --- | --- | --- | --- | --- | --- |
| $1 \times 10^{-3}$ | $1.02 \times 10^{-3}$ | $0.97 \times 10^{-3}$ | $1.04 \times 10^{-3}$ | $1.05 \times 10^{-3}$ | $1.00 \times 10^{-3}$ | $1.00 \times 10^{-3}$ |
| $1 \times 10^{-4}$ | $1.09 \times 10^{-4}$ | $0.91 \times 10^{-4}$ | $1.06 \times 10^{-4}$ | $0.99 \times 10^{-4}$ | $1.02 \times 10^{-4}$ | $1.00 \times 10^{-4}$ |
| $1 \times 10^{-5}$ | $1.10 \times 10^{-5}$ | $0.95 \times 10^{-5}$ | $1.00 \times 10^{-5}$ | $0.90 \times 10^{-5}$ | $1.10 \times 10^{-5}$ | $1.00 \times 10^{-5}$ |
| $2.8 \times 10^{-6}$ | $4.00 \times 10^{-6}$ | $3.00 \times 10^{-6}$ | $2.50 \times 10^{-6}$ | $2.00 \times 10^{-6}$ | $3.00 \times 10^{-6}$ | $3.00 \times 10^{-6}$ |

### Power Simulation

To evaluate the power of the proposed method, we conducted extensive simulations using the same gene *EP B*41. We simulate 10^4^ summary statistics from *N* (***A** × Δ, **R***) where ***A*** denotes the signs of associations which determine the risk or protective effects of causal variants, *Δ* denotes different settings which determine the effect sizes of causal variants, and ***R*** is the corresponding LD matrix of the gene *EP B*41. We assume that there are 3 causal SNPs. The effects of the three causal SNPs are randomly set by drawing 3 elements of ***A*** equal to 1 or −1, and setting the effect of the rest of SNPs in the gene as 0. Table 2 shows the estimated power at 2.80 *×* 10^*−*6^ significance level under three combinations of ***A*** for different settings of *Δ*: a set of fixed values of *Δ* and two randomly simulated *Δ*, one from uniform distribution, and the other from normal distribution. Overall, the proposed OWC test performs robustly across all scenarios and has the best performance compared to the other five tests, which may be attributed to the fact that our proposed method includes two kinds of burden tests and two kinds of quadratic tests, and ultimately derives an optimal test incorporating the four classes of methods. Thus, the OWC test reaches the maximal power. From the first three settings of three fixed values of *Δ*, we can clearly see that the power of the S2T and GATES methods increases as the effect size increases, which implies their performance may suer when there are weak eects, compared to the other methods. Because S2T is a quadratic method and GATES is a p-value based method, they are all robust to the direction of effects among causal SNPs. The results of the last three settings from the same normal distribution of *Δ* also verify this conclusion. Comparing the results of the middle three settings from a uniform distribution of *Δ*, the burden test ST has the worst performance when there are different directions of effects. Even though the adaptive methods AT and aSPU all consider the combination of the burden test and quadratic test methods, they also suffer a relatively small loss of power when weak effect size and different directions of effects are considered. Our proposed test OWC is robust in the presence of different directions as well as weak effect sizes.

**Table 2:** Estimated power (%) under 2.8 *×* 10^*−*6^ significance level for different tests. Data are simulated from *N*(***A** × Δ, **R***). ***A*** has 3 nonzero elements with different signs.

| nonzero $\Delta$ | nonzero $\mathbf{A}$ | AT | S2T | ST | GATES | aSPU | OWC |
| --- | --- | --- | --- | --- | --- | --- | --- |
| (4,2,1) | (1,1,1) | 68.0 | 8.5 | 68.5 | 16.5 | 83.5 | 95.0 |
| (4,4,2) | (1,1,-1) | 69.0 | 45.0 | 45.0 | 28.5 | 65.6 | 93.5 |
| (2,5,4) | (1,-1,-1) | 92.5 | 74.5 | 76.0 | 65.5 | 62.0 | 99.5 |
| U(1,5) | (1,1,1) | 86.0 | 37.0 | 87.0 | 24.0 | 86.0 | 97.0 |
| U(2,6) | (1,1,-1) | 71.5 | 71.5 | 19.0 | 58.0 | 68.0 | 85.5 |
| U(2,6) | (1,-1,-1) | 70.0 | 70.5 | 19.5 | 65.5 | 64.5 | 88.5 |
| N(3,4) | (1,1,1) | 83.0 | 60.0 | 83.5 | 74.5 | 88.5 | 94.5 |
| N(3,4) | (1,1,-1) | 68.0 | 70.0 | 33.0 | 72.5 | 69.5 | 82.5 |
| N(3,4) | (1,-1,-1) | 66.0 | 61.5 | 33.0 | 78.0 | 73.5 | 87.0 |

### Application to the Schizophrenia GWAS Summary Data

We applied the proposed method OWC and the other five tests to two schizophrenia (SCZ) summary datasets, which were downloaded from the Psychiatric Genomics Consortium (PGC) website (see URL https://www.med.unc.edu/pgc/results-and-downloads/): a meta analysis of SCZ GWAS dataset with 13,833 cases and 18,310 controls, denoted as SCZ1 (Ripke et al., 2013), and a more recent and larger dataset with 36,989 cases and 113,075 controls, denoted as SCZ2 (Schizophrenia Working Group, 2014). The summary data consists of MAF, estimated effect size, odds ratio, and p-value for 560,833 SNPs on 17,866 genes in SCZ1 and 557,511 SNPs on 17,824 genes in SCZ2. We define a gene including all of the SNPs from 20 kb upstream to 20 kb downstream of the gene by following Wu et al. (2010) and test the association between the gene and the trait using OWC and other five tests. We use the 1000 Genomes project reference panel (1000 Genomes Project Consortium, 2012) to compute the LD for SNPs within each gene contained in the two datasets. In order to make fair comparsions among the six tests, we remove SNPs with MAF*<* 0.05 as well as one of a pair of SNPs with pairwise LD *r*^2^ *>* 0.5. After SNPs pruning, 174,648 SNPs on 17,467 genes in SCZ1 data and 174,275 SNPs on 17,420 genes in SCZ2 data remained in our final analysis. P-values of our method OWC and aSPU are estimated by performing 10^6^ permutations. The genome-wide significance level for the gene-based genome-wide association test is *≈* 2.80 *×* 10^*−*6^, which is the Bonferroni corrected significance level. First, we applied our method and the other comparable methods to the SCZ1 data (Ripke et al., 2013) of 20,899 individuals to identify SCZ-associated genes. We then searched for genome-wide significant genes in the larger SCZ2 dataset (Schizophrenia Working Group, 2014) of 150,064 individuals for partial validation. Figure 1 shows a Venn diagram of the number of significant genes identified in SCZ1 by our proposed method OWC compared to aSPU, GATES, and Guo and Wu’s three methods (denoted as GW, which represents the aggregate of genes identified by S2T, ST and AT combined). OWC identified 37 significant genes, aSPU identified 21 significant genes, GATES identified 17 significant genes, and GW identified 23 significant genes in total. Among these 37 significant genes identified by OWC, 13 (around 35%) and 21 (around 57%) contained the genome-wide significant SNPs (p-value*<* 5 *×* 10^*−*8^) within 20 kb in the SCZ1 data and the SCZ2 data, respectively, offering significant validation of the identified genes. Clearly, our method identified more associated genes than the other methods.

**Figure 1.**
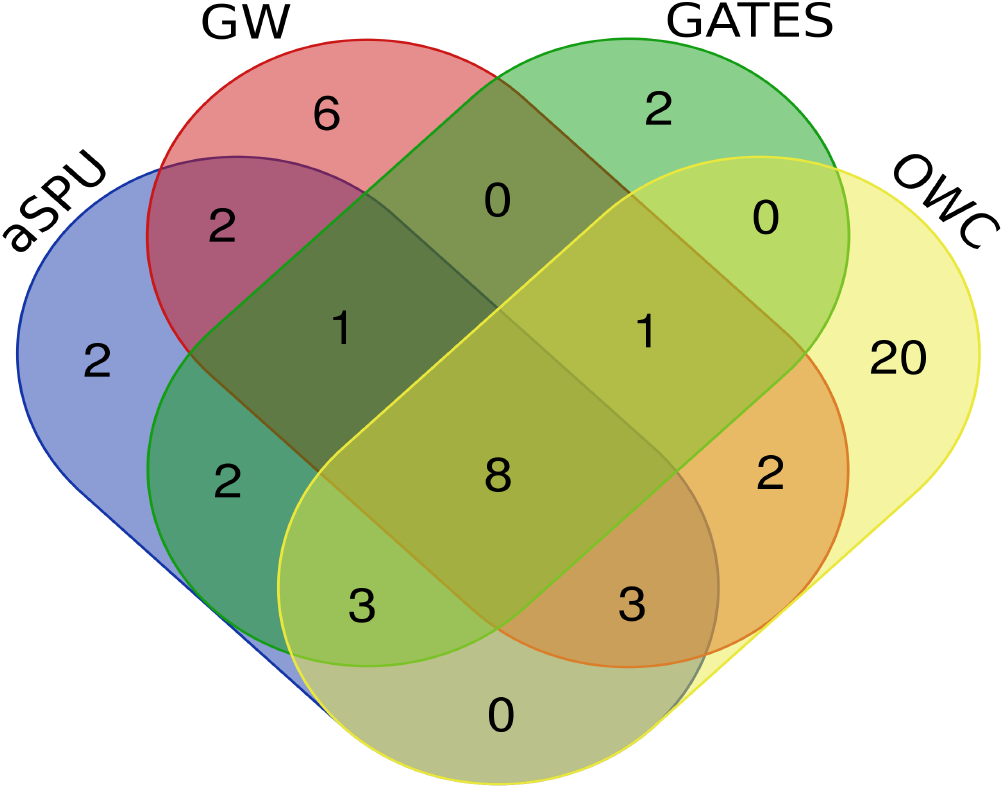
Venn diagram of the number of significant genes identified by OWC, aSPU, GATES, and GW for SCZ1.

Overall, a total of 52 significant (and unique) genes were identified in the SCZ1 data across all four tests. Supplementary Table S1 shows the significant genes, with associated p-value and minimum p-value, identified by OWC, aSPU, GATES, or GW in SCZ1 and SCZ2, respectively. Next, we applied the four methods to the SCZ2 data in which OWC identified 135 significant genes, aSPU identified 110 significant genes, GATES identified 116 genes, and GW identified 125 significant genes in total. As expected, because the sample size of the SCZ2 dataset is much larger than that of SCZ1, more genes were identified as significant by all of the methods. The number of significant genes identified by OWC compared to aSPU, GATES and GW can be seen in Figure 2. Again, our method is more powerful than the other methods in terms of the number of significant genes identified. However, each test identified some unique genes missed by the others, highlighting that different tests may be more powerful in different scenarios. Combined, the four methods identified 188 significant and unique genes in the SCZ2 data (Supplementary Table S2).

**Figure 2.**
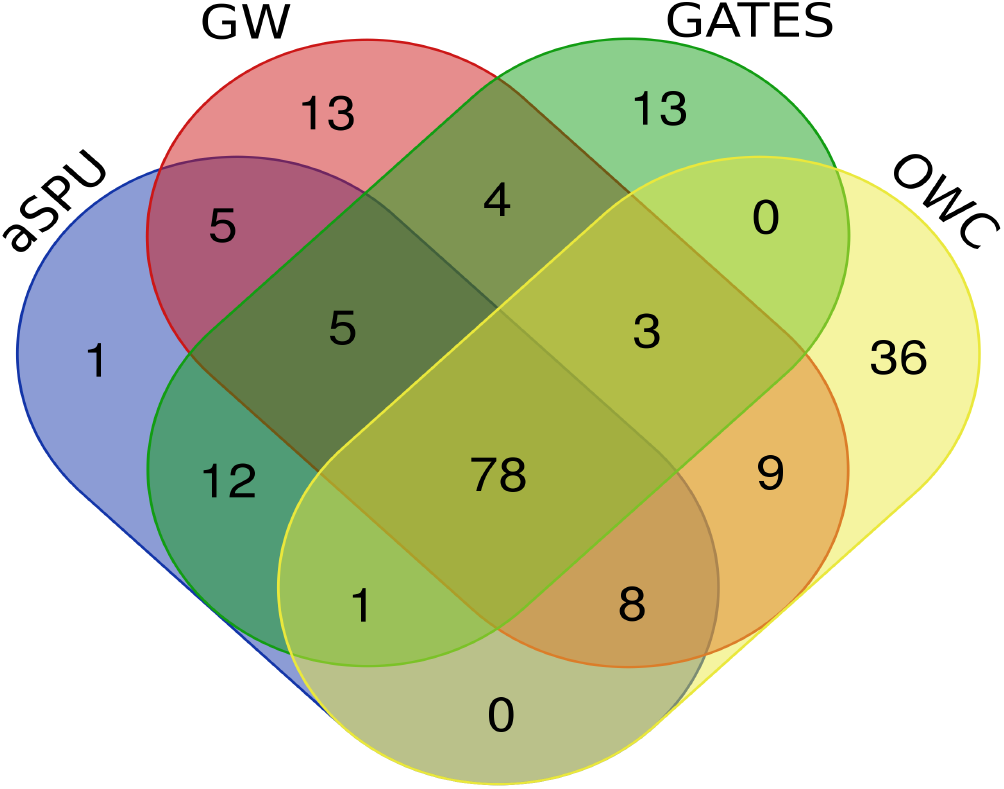
Venn diagram of the number of significant genes identified by OWC, aSPU, GATES, and GW for SCZ2.

### Application to the T2D GWAS Summary Data

We also conducted a comprehensive analysis of fasting glucose GWAS summary data for type 2 diabetes (T2D) from the UK Biobank component of the European DIAMANTE study (denoted as UKB), which included over 440,000 individuals of European ancestry with 19,119 cases and 423,698 controls. The analysis for the dataset was restricted to HRC variants and was conducted using the UK Biobank Resource under Application Number 9161 (McCarthy). The GWAS summary data can be downloaded from http://www.type2diabetesgenetics.org/informational/data. Similar to the SCZ dataset, the UKB summary data also consists of MAF, effect size estimate, odds ratio, and p-value for approximately 17,850 genes. We applied the same procedure used to filter and analyze the SCZ data on the UKB data. We used 0.05*/*17, 850 *≈* 2.80 *×* 10^*−*6^ as the significance level and performed 10^6^ permutations for our proposed method OWC and aSPU.

Figure 3 shows the Venn diagram comparing the number of significant genes identified by our proposed method compared with aSPU, GATES and GW: OWC identified 133 significant genes, whereas aSPU, GATES, and GW identified 99, 94, and 95 significant genes, respectively. Among these 133 significant genes identified by OWC, 66 (around 50%) contained the genome-wide significant SNPs (p-value *<* 5 *×* 10^*−*8^) within 20 kb in the UKB data. Based on the UKB analysis, we can further conclude that our OWC method performed the best compared to the other test methods in terms of the number of significant genes identified. The 155 significant and unique genes identified by all four methods can be seen in Supplementary Table S3.

**Figure 3.**
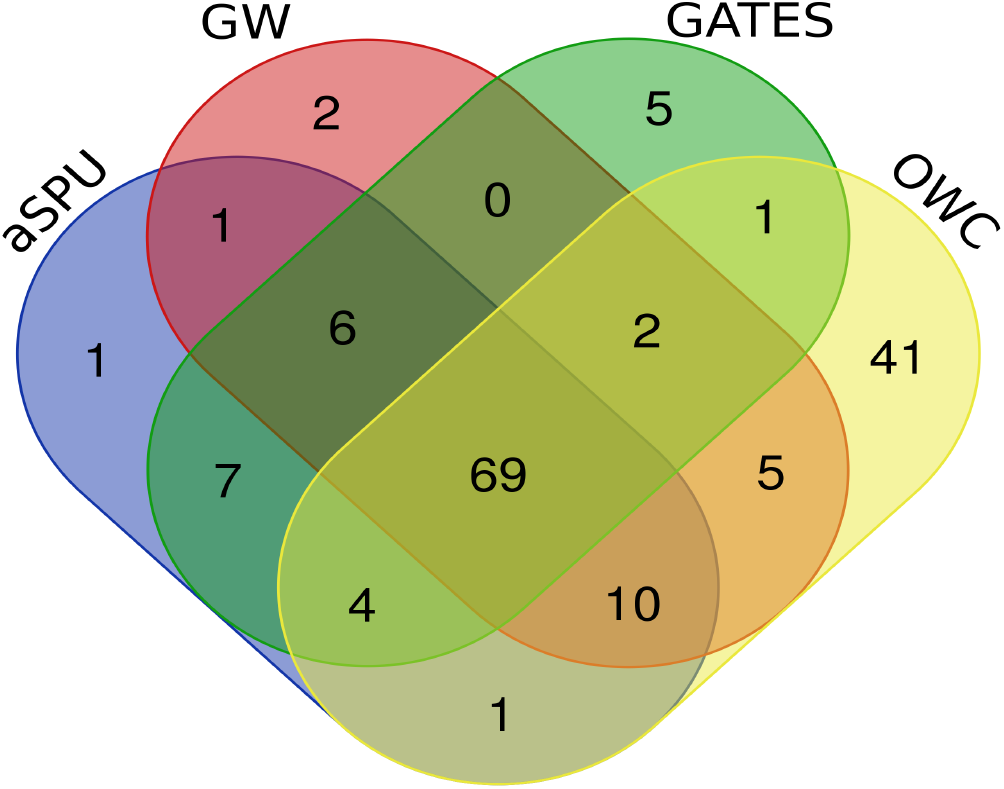
Venn diagram of the number of significant genes identified by OWC, aSPU, GATES, and GW for UKB.

### Summary of Real Data Analaysis

Several genes that contain highly significant SNPs in the SCZ and UKB datasets were identified by all four methods, giving credence to the power and validity of OWC. Many of the genes identified as significant also have important biological implications regarding the etiology of the associated disease. For example, all four methods identified several genes in the SCZ data which encode for micro RNA (miRNA), such as MIR137/MIR2682: Altered miRNA expression has been considered a plausible mechanism for development of schizophrenia and other psychiatric disorders such as bipolar disorder (BD) and major depression (MD) (Duan et al., 2014; Maffioletti et al., 2014; Kwon et al., 2013). Likewise, *NU BP L* is a gene involved with respiratory complex I, a mitochondrial enzyme which may also be associated with the development of schizophrenia (Calvo et al., 2010; Karry et al., 2004; Ben-Shachar, 2017). In the UKB dataset, all four methods identified *V EGF B* as significant, a gene which has been considered a potential target in T2D therapy due to its elusive role in modulating insulin resistance (Hagberg et al., 2012), as well as *RCAN*2 and *MT RNR*2*L*1, both of which have been implicated in other studies (Xin et al., 2016; Veilleux et al, 2015).

The OWC method also identified biologically relevant genes that were missed by the other methods. For instance, OWC was the only method to identify *SGT A* as significant in the UKB dataset. *SGT A* plays a role in poly-cystic ovary syndrome (PCOS) in women, an endocrine disease involving insulin resistance and co-morbidity with T2D (Goodarzi et al., 2011), indicating that this gene may also be involved with T2D pathogenesis.

Additionally, only OWC identified gene *GRM*7 in the SCZ data as significant. This gene produces a type of glutamate receptor, dysfunction of which has long been implicated in schizophrenia (Ohtsuki et al., 2008; Li et al., 2016), and studies regarding individualized therapy have considered *GRM*7 as a potential biomarker for risperidone response (Sacchetti, et al., 2017). The p-value for most significant SNP in *GRM* 7 was 1.12E-06 for both SCZ1 and SCZ2, highlighting OWC’s ability to identify weakly associated variants, which are often overlooked by traditional methods.

## Discussion

The weighting of genetic variants plays an important role in boosting the power of a test in genetic association studies. In this paper, we propose a novel gene based genetic association test, the optimal weighted combination (OWC) test, which is a general linear weighted test statistic which incorporats four different weighting schemes (two constant weights and two weights proportional to normal statistic ***Z***). Four well-known methods: burden test, WSS, SSU, and the score test are included in OWC as special cases. If we focus on the summary data obtained from rare variants analysis, we can set the elements on the diagonal of matrix ***A*** as the beta distribution density function with pre-specified shape parameters as 1 and 25 evaluated at the corresponding sample MAF in the data. In this situation, the method SSU contained in our method OWC becomes the SKAT method. Thus, both the SKAT and SKAT-O methods can additionally be viewed as special cases of our method. If we focus on data from transcriptome-wide association studies, we can set the weights of WSS and the elements of the diagonal of matrix ***A*** as the corresponding estimated cis-effects on gene expression. Then, the two methods WSS and SSU contained in our method become PathSPU(1) and PathSPU(2) method proposed by Wu and Pan (2018). Therefore, the OWC is an optimally weighted combination test that can reach maximized power.

We analytically derive that a score test based on the aggregated variants in a gene is the same as a score test based on the weighted combination of Z-scores of SNP based association tests in the gene. Additionally, we show that the general optimal weight can be obtained when the general linear weighted test statistic reaches its maximum. For example, the score test, a special case of OWC, reaches its maximum when we consider the correlation matrix ***R*** among SNPs. When we ignore the correlation among SNPs (it means that we set ***R*** = ***I***), OWC becomes the SSU test, which reaches its maximum. Through simulation studies and real data analyses, we demonstrate that the proposed optimal test often outperforms other comparison methods such as ST, S2T, AT, aSPU, and GATES, which are some of the most popular methods based on summary data.

When we perform association tests based on a large number of genes in whole-genome sequencing studies, different disease models may exist: some of the models are likely to include many causal variants whose effects are in the same directions while other models may include few causal variants or the causal variants whose effects are in different directions; or some of the models are likely to include many weakly associated variants while others may include a few of strongly associated variants. Because the true disease model is usually unknown, there is no uniformly most powerful test to detect single trait associated genes; an association test may perform well for one dataset, but not necessarily for another. For example, both schizophrenia and type 2 diabetes are representative of complex diseases with common, often weakly associated variants in which many of these variants may be working in tandem to produce the disease. A robust, flexible method such as OWC may better elucidate these weakly associated variants so that their role in disease etiology can be understood further. Indeed, the proposed OWC can be an attractive tool for many situations, because it adapts to the underlying biological disease model by selecting ***ρ*** based on the data.

Our proposed method provides an alternative approach with extreme scalability and robust performance. This method only needs the publicly available GWAS summary statistics as input, without the need to access raw genotype and phenotype data. We expect that researchers will be able to identify novel disease associated genes by employing the proposed method to analyze publicly available GWAS summary data and shed more light onto underlying mechanism of disease. In this paper, we focus on the application of our proposed method using single trait GWAS summary data, however OWC can be easily extended to multiple traits GWAS summary data. We have implemented the proposed method in an R program which is publicly available online at github.

## Appendix

Use the notation in the Method section. Consider a raw data set of a sample including *n* individuals, where each individual has been genotyped at M variants in a genomic region (gene or pathway). Denote *y*_*i*_ as the trait value of the *i^th^* individual for either a quantitative or qualitative trait (1 for cases and 0 for controls for a qualitative trait) and denote *X*_*i*_ = (*x_i1_*, …, *x*_*iM*_)′ as the genotypic score of the *i^th^* individual, where *x_im_ ∈ {*0, 1, 2*}* is the number of minor alleles that the *i^th^* individual has at the *m^th^* variant.

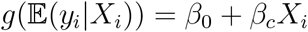

where *g*(*⋅*) is a monotone “link” function and *β_c_* is the parameter of interest. Testing the association of the genetic variants in the considered region is equivlent testing the effect of the weighted combination of variants 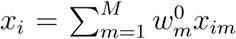. Under the generalized linear model, we can use the score test statsitic to test the null hypothesis *H*_0_ : *β_c_* = 0, which is given by

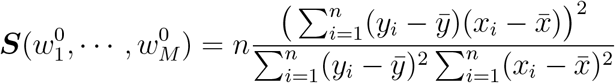

where the score test statistic ***S*** can be viewed as a function of weight 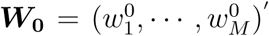. Let ***X*** = (*X*_1_, …, *X*_*n*_)′, *Y* = (*y*_1_, …, *y*_*n*_)′ and 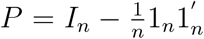 where 1_*n*_ represents a column vector containing all ones. Then, we have 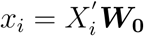. We can rewrite the score test as:

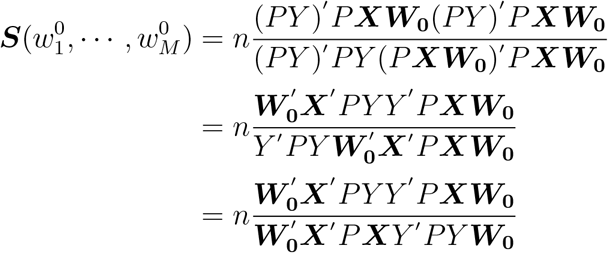

where the second equation holds because we have *P* = *P*′ and *PP*′ = *P*, and the third equation holds because *Y*′*P Y* is a constant.

To test the association between a single trait and a single variant, we usually employ a Z test. To test the main effect of the *m^th^* variant in the considered region, we use the Z test: 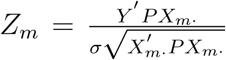 where 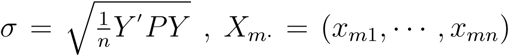. Let LD matrix ***R*** = *diag*(***D***)^*−*1*/*2^***D**diag*(***D***)^*−*1*/*2^ where ***D*** = ***X***′*P**X*** and *diag*(***D***) denote the diagonal matrix of ***D***. When GWAS summary statistics such as the Z-statistics and the LD matrix for SNP-SNP correlations are available, the score test can be written as:

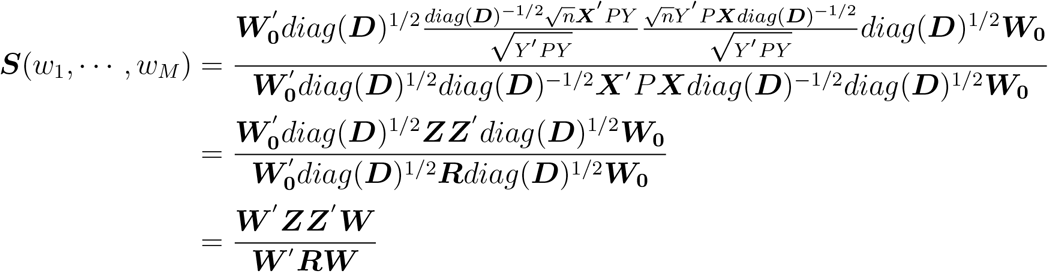

where ***Z*** = (*Z*_1_, …, *Z*_*M*_)′ and ***W*** = (*w*_1_, …, *w*_*M*_)′ = *diag*(***D***)^1*/*2^***W***_0_. From Equation (1), the score test statistic ***S*** is equivalent to a linear weighted test statistic based on Z-scores:

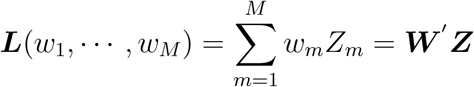

where ***Z*** follows multivariate normal distribution with mean **0** and covariance matrix ***R*** under null hypothesis (Zhang et al. 2018). This conclusion clearly demonstrates that testing the weighted combination of variants in a considered region using the score test is the same as using the weighted combination of Z-scores for the variants in the considered region.

## Supplemental Data

Supplemental Data includes three tables: Supplementary Table S1: the OWC and the other comparison methods are applied to the SCZ1 data and significant genes identified are reported in Supplementary Table S1; Supplementary Table S2: the OWC and the other comparison methods are applied to the SCZ2 data and significant genes identified are reported in Supplementary Table S3; Supplementary Table S3: the OWC and the other comparison methods are applied to the UKB data and significant genes identified are reported in Supplementary Table S3.

## Acknowledgments

X. Raymond Gao was supported by National Institutes of Health (NIH; Bethesda, MD, USA) grants R01EY027315 and RF1AG060472. X Wang was supported by the University of North Texas Foundation which was contributed by Dr. Linda Truitt Creagh. The content is solely the responsibility of the authors and does not necessarily represent the views of the University of North Texas Foundation and Dr. Linda Truitt Creagh.

